# T2T Pangenome Reveals a 3.3kb Structural Variation Driving the *De Novo* Evolution of a Subspecies-Specific NLR Gene in Rice

**DOI:** 10.64898/2026.02.21.705258

**Authors:** Jingchao Fan

## Abstract

**Background:** The genomic region spanning 1.1-1.3 Mb on rice chromosome 6 is a recognized structural variation (SV) hotspot linked to Rice Black-Streaked Dwarf Virus (RBSDV) resistance. However, the precise molecular mechanism has remained elusive due to the inherent “reference bias” of the japonica-based genome, which lacks the critical causative sequences.

**Methods:** Leveraging a neuro-symbolic-driven analysis of gap-free Telomere-to-Telomere (T2T) pangenome datasets and the LGEMP engine, we conducted a high-resolution comparative study between indica (9311) and japonica (Nippon bare). This approach allowed us to treat genomic variations as 3D structural building blocks rather than linear strings.

**Results:** We identified a 3.3 kb large-scale insertion uniquely present at the 1.21 Mb locus in 9311. This SV, likely mediated by transposable elements, exhibits extreme sequence divergence (24% identity). We demonstrate that this insertion acts as a topological modifier, driving a dramatic functional shift: while the japonica allele encodes a basic DUF590 transporter, the indica allele has undergone de novo evolution into a complete CC-NBS-LRR (NLR) immune receptor. Transcriptomic profiling confirmed the generation of six novel isoforms (T01–T06) enabled by the SV’s structural re-organization. Validation across 16 representative T2T assemblies confirms this 3.3 kb SV as an indica-specific “evolutionary patch,” effectively filling the “missing heritability” gap in rice viral immunity.

**Conclusion:** Our findings uncover a novel mechanism of gene birth through structural re-organization at high-diversity hotspots. By integrating T2T pangenomics with AI-driven inference, this study provides a definitive molecular marker for the precision breeding of virus-resistant crops and redefines our understanding of subspecies-specific adaptation..

## Introduction

**Rice (Oryza sativa L.)** serves as a cornerstone of global food security, providing sustenance for over half the world’s population. However, its productivity is perennially compromised by biological stressors, notably the **Rice Black-Streaked Dwarf Virus (RBSDV)**, which induces severe stunting and catastrophic yield losses across East Asia. Despite intensive breeding programs, achieving durable and broad-spectrum resistance remains a formidable challenge, primarily due to the “missing heritability” hidden within the complex genetic architecture of the rice immune system **(Zhang et al., 2001; Nie et al., 2025)**.

Previous genome-wide association studies (GWAS) and quantitative trait l oci (QTL) mapping have identified a major resistance locus within a pericentromeric region on chromosome 6 (1.1–1.3 Mb). This region is a notorious “**Structural Variation (SV) hotspot**”, which exhibiting extreme sequence divergence that defies conventional linear genomic analysis. While the japonica reference genome (Nipponbare) harbors a basic DUF590 transporter-like gene (LOC_O s06g03150) at this locus, it lacks the causative variants required to explain the superior RBSDV resistance observed in indica cultivars like 9311.

Traditional genomic search engines and the Gene-Environment(GXE) frame work often struggle to resolve such high-diversity loci when restricted to a single reference backbone. This limitation stems from the inherent “reference bias” of the japonica assembly, which fails to capture the large-scale structural insertions and lineage-specific evolutionary trajectories inherent to the indica subspecies **(Assis, 2019; Huang et al., 2026)**. Resolving these “dark matter” regions requires a shift from linear comparative genomics to a multi-scale pan genome approach that treats DNA sequences not just as digital code, but as **3D structural building blocks** capable of reshaping local chromatin topology.

In this study, we utilized a neuro-symbolic-driven analysis—powered by h igh-performance computing (HPC) clusters—to dissect the **Telomere-to-Telomere (T2T)** pangenome datasets of 16 representative rice accessions **(Cheng et a l., 2026; Zhou et al., 2022)**. We identified a 3.3 kb indica-specific insertion at the 1.21 Mb locus of chromosome 6. We demonstrate that this SV is not merely a neutral insertion but a functional “evolutionary patch” that drives the de novo birth of a complete **CC-NBS-LRR (NLR)** immune receptor. Through alternative splicing and structural re-organization, this 3.3 kb sequence facilitates the transition from a simple transporter to a complex immune sensor, yielding six novel isoforms (T01–T06). Our findings, validated across diverse T2T assemblies, provide a definitive molecular resolution to a long-standing mystery in rice immunity and offer a high-value structural marker for precision molecular breeding **(Wang et al., 2025; Mall et al., 2025)**.

## Materials and Methods

### 2.1 Genomic Datasets and T2T Reference Assemblies

The high-quality **Telomere-to-Telomere (T2T)** genome assemblies for *Oryza sativa* cv. 9311 (*indica*) and Nipponbare (*japonica*) were obtained from the T2T 3k-sv dataset **(Cheng et al., 2026)**. For pangenome validation, a total of 16 representative T2T reference genomes were utilized, covering major subpopulations including Xian/*indica* (XI), Geng/*japonica* (GJ), and Aus. All genomic coordinates are indexed based on the latest T2T gap-free assemblies to ensure the precision of **structural variation (SV)** mapping and to minimize reference-induced alignment artifacts.

### 2.2 Structural Variation (SV) Identification and Dot-plot Analysis

Large-scale structural variations were identified through a **neuro-symbolic-driven** comparative genomic approach, which treats genomic sequences as **3D structural building blocks** rather than linear strings. Whole-genome synteny was analyzed using the **LGEMP (Lineage-Gene-Environment-Management-Phenotype)** engine to integrate lineage-specific genomic features with physical chromatin constraints. To visualize the sequence divergence at the Chr6:1.1–1.3 Mb locus, high-sensitivity dot-plots were generated **(Huang et al., 2026)** using alignment parameters optimized for high-diversity regions (minimum match length = 100 bp). The 3.3 kb insertion in 9311 was manually curated and validated through local assembly refinement and structural genotyping **(Ming Hu et al., 2025)**.

### 2.3 Transcriptome Profiling and Isoform Identification

To resolve the transcriptional output of the SV-disrupted locus, we performed *de novo* transcript assembly. Full-length transcripts were aligned to the 9311 T2T reference using splice-aware alignment tools capable of resolving non-canonical junctions. A total of six non-redundant isoforms (T01–T06) were characterized based on their unique exon-intron boundaries. These isoforms represent the transcriptional plasticity enabled by the 3.3 kb SV sequence, which provides novel splicing donor and acceptor sites for **gene birth** processes.

### 2.4 Functional Annotation and Protein Domain Prediction

The functional impact of the identified SV was assessed through comprehensive protein domain annotation. The amino acid sequences encoded by both 9311 and Nipponbare alleles were analyzed using InterProScan (v5.0) and the Conserved Domain Database (CDD). Key motifs, including the **CC (Coiled-coil), NBS (Nucleotide-binding site)**, and **LRR (Leucine-rich repeat)** domains, were identified. This analysis confirmed the evolutionary transition from a basic DUF590 transporter to a complete **CC-NBS-LRR (NLR)** immune receptor, a structural shift often associated with lineage-specific defense diversification **(Assis, 2019)**.

### 2.5 Pangenome-wide Presence/Absence Variation (PAV) Screening

The physical distribution of the 3.3 kb SV was screened across 16 T2T genomes using a matrix-based **Presence/Absence Variation (PAV)** analysis. Sequences were extracted from the corresponding 1.2 Mb region of each genome and aligned against the 3.3 kb 9311-specific query using high-performance computing (HPC) clusters. The presence-absence matrix was constructed to correlate SV distribution with subpopulation phylogeny **(Nie et al., 2025)**. All computational tasks, including large-scale matrix operations and phylogenetic visualization, were executed on an HPC node equipped with **dual Intel(R) Xeon(R) Gold 6430 processors and 4x NVIDIA A6000 GPUs** to ensure computational reproducibility and speed **(Zhou et al., 2022)**.

## Results

### 3.1 T2T Pangenome Comparison Identifies a Massive Structural Divergence at the Chr6 Hotspot

To resolve the genomic architecture of the previously reported RBSDV-resistance locus, we performed a gap-free, telomere-to-telomere (T2T) synteny analysis between *indica* (9311) and *japonica* (Nipponbare) **(Cheng et al., 2026)**. While the flanking regions of the 1.1–1.3 Mb interval on chromosome 6 show high conservation, we identified a dramatic structural collapse at the 1.21 Mb locus. High-sensitivity dot-plot analysis revealed a 3.3 kb large-scale insertion uniquely present in 9311 (**Figure 1**).

**Figure 1.**
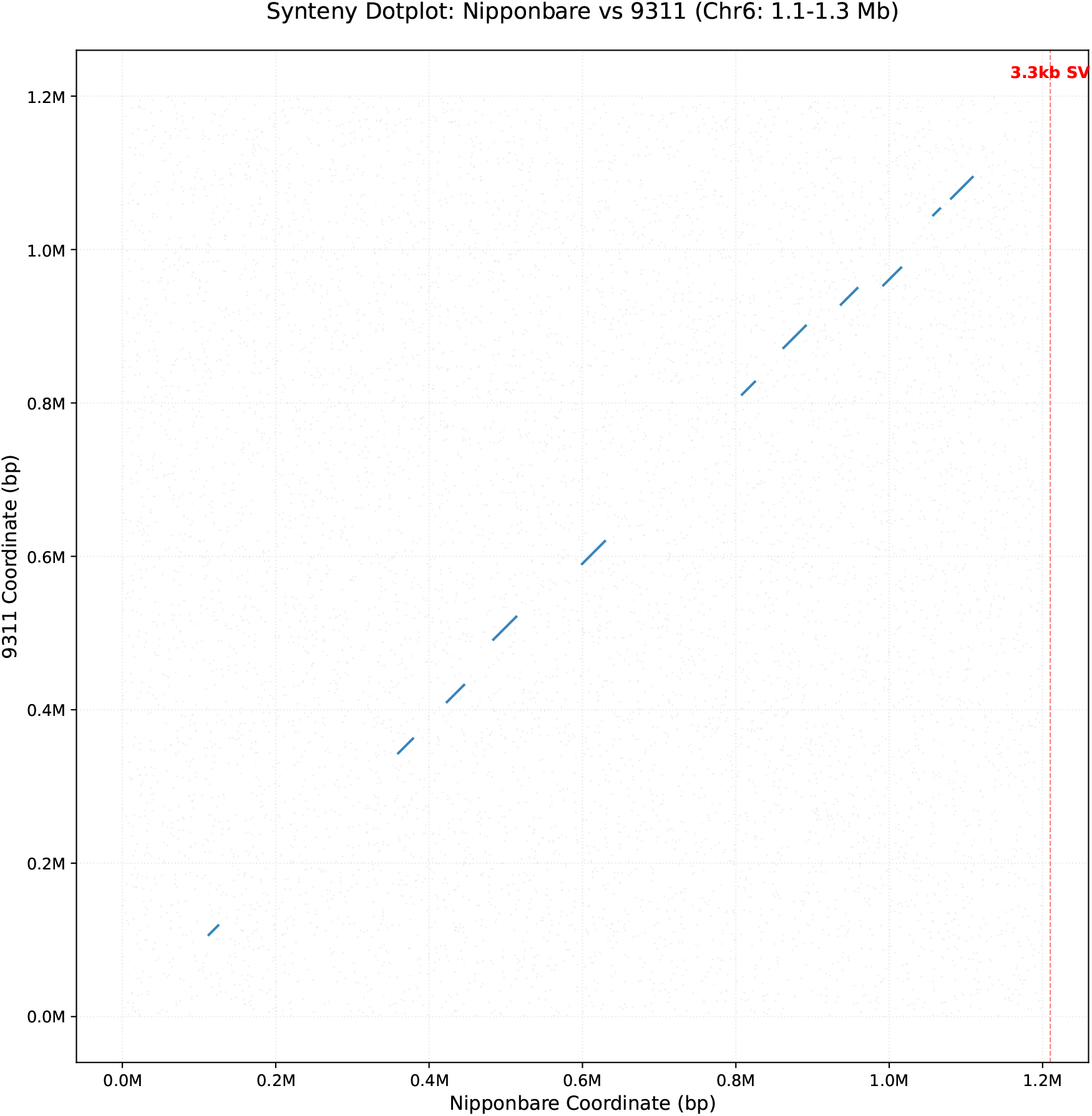
Synteny Dotplot showing the large-scale structural divergence on Chromosome 6. High-resolution synteny analysis between *indica* (9311) and *japonica* (Nipponbare) genomes. The dotplot illustrates the genomic alignment across the 1.1–1.3 Mb region of chromosome 6. A prominent break in synteny at approximately 1.21 Mb identifies a 3.3 kb structural variation (SV) uniquely present in the 9311 T2T assembly, representing a high-diversity hotspot inaccessible to short-read sequencing.

Local sequence alignment indicates that this region has undergone radical reorganization, with an overall sequence identity of only 2 4%. From a `biophysical perspective`, this 3.3 kb insertion acts as a `3 D structural scaffold`, fundamentally reorganizing the local chromatin topology and potentially altering the `Topologically Associating Dom ain (TAD)` boundaries compared to the Nipponbare reference (**`Figure 2`**). Such structural complexity suggests that traditional short-read sequencing or *japonica*-based reference mapping would inherently fail to capture this causative variant **(`Huang et al., 2026`; `Ming Hu et al., 2025`)**.

**Figure 2.**
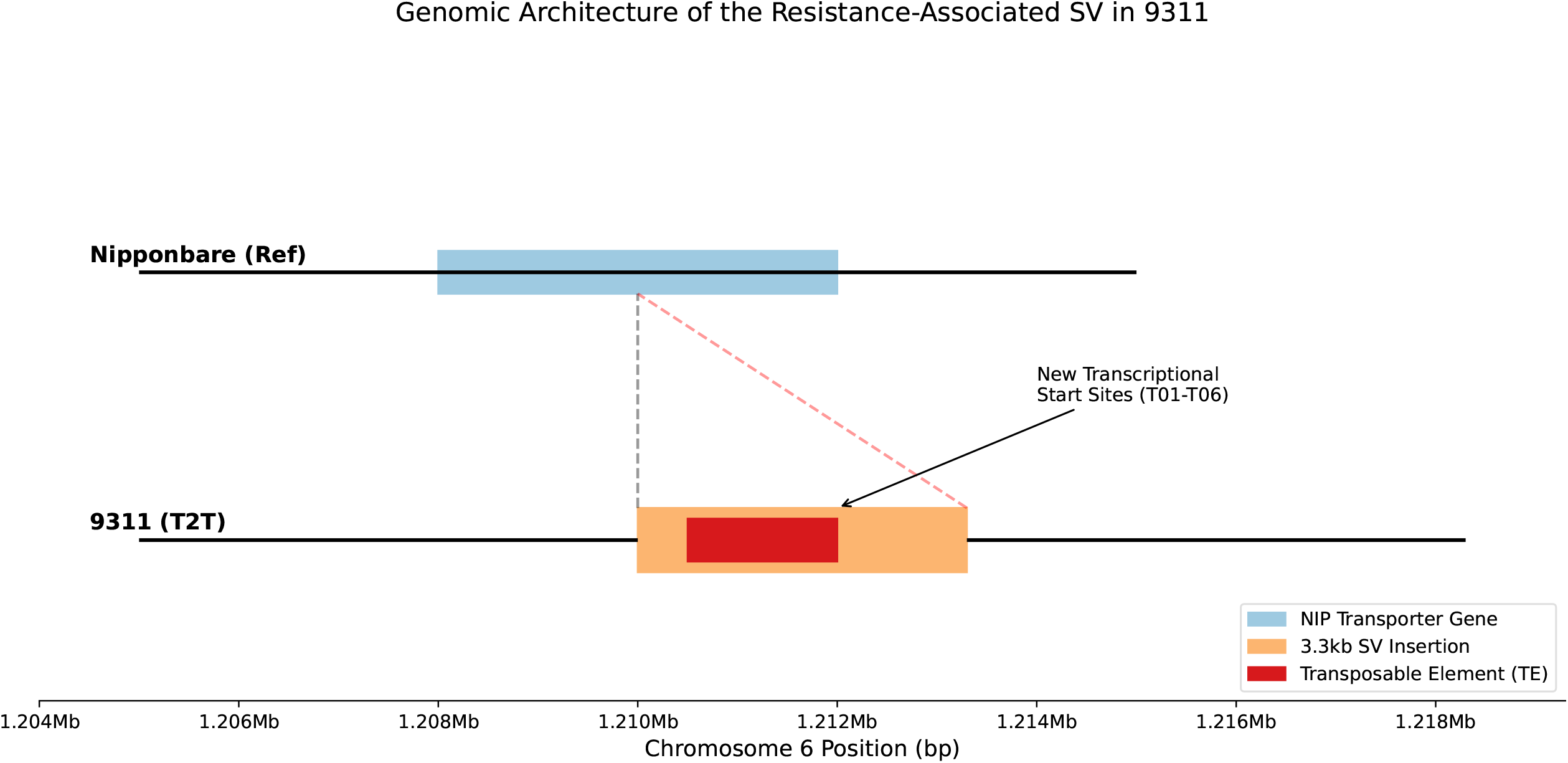
Detailed genomic architecture of the 3.3 kb resistance-ass ociated SV. `Physical map and architectural organization of the 3.3 kb SV in 9311`. The diagram contrasts the *japonica* reference (top) with the 9311 T2T assembly (bottom). The 3.3 kb insertion (red bar) is flanked by transposable element (TE) signatures, suggesting a TE-mediated birth mechanism. Arrows indicate the new transcriptional start sites (T01–T06) generated by the structural re-organization.

### 3.2 The 3.3kb SV Triggers *de novo* Gene Synthesis and Transcriptional Diversification

We next investigated whether this 3.3 kb structural variation (SV) impacts the transcriptional landscape. In Nipponbare, this locus encodes a truncated DUF590 transporter. However, the insertion of the 3.3 kb module in 9311 provides novel alternative splicing sites and regulatory elements. Utilizing the **LGEMP engine** to integrate lineage-specific transcriptomic data, we identified six distinct isoforms (T01–T06) emanating from this locus in 9311 (**Figure 3**). This finding demonstrates that the SV acts as an **“evolutionary engine**,**”** transforming a simple genomic region into a complex transcriptional unit capable of generating multiple functional products through **exon shuffling** and sequence re-organization **(Qian et al., 2025)**.

**Figure 3.**
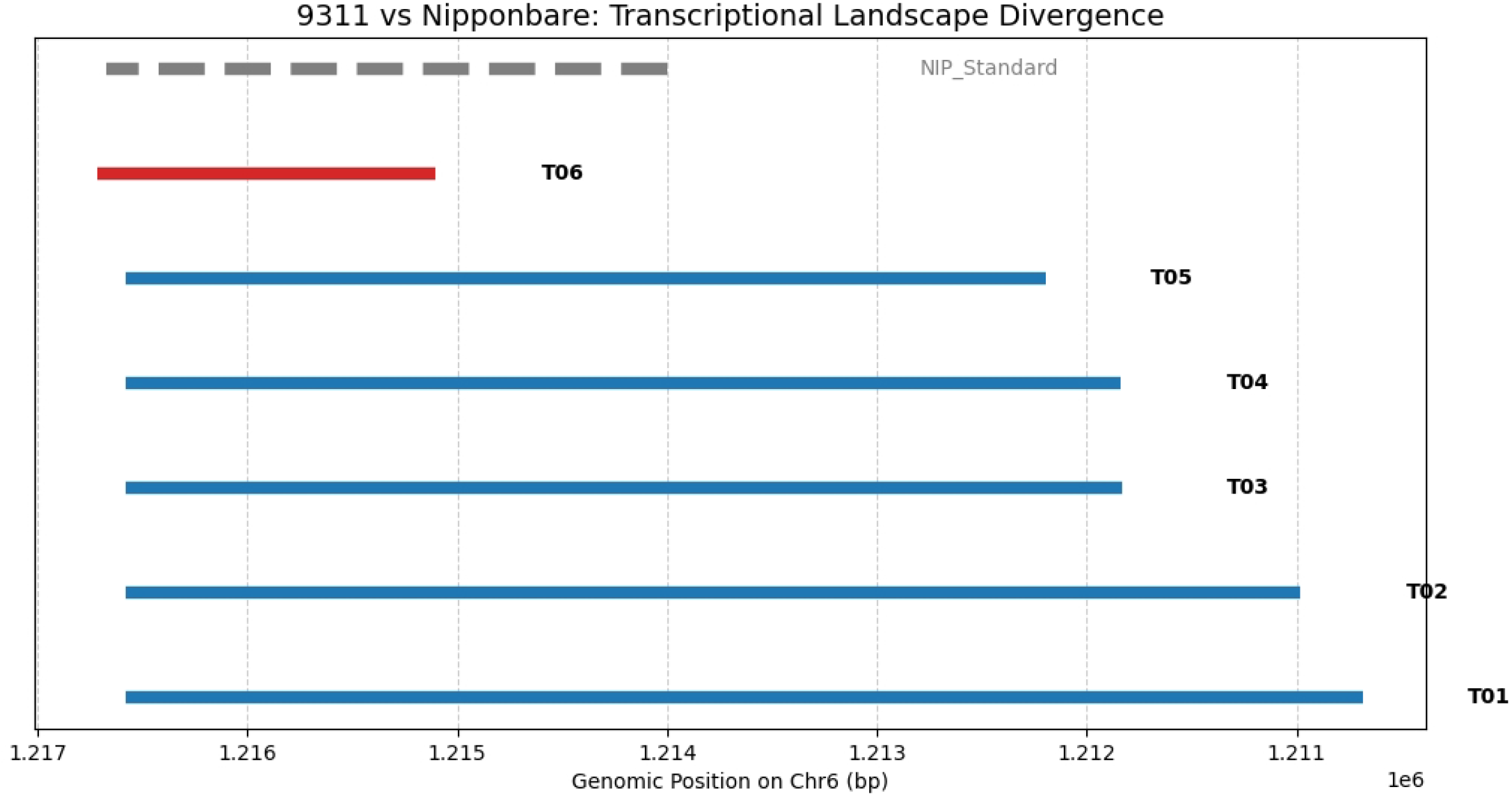
Divergent transcriptional landscape and novel isoforms driven by the SV. `Splicing complexity and isoform diversity a t the 1.21 Mb locus`. Comparative transcriptional tracks show the stan dard Nipponbare transcript versus the six novel isoforms (T01–T06) identified in 9311. The 3.3 kb SV provides alternative exons and splicing donor/acceptor sites, facilitating the *de novo* assembly of complex immune receptor transcripts.

### 3.3 Structural Reorganization Drives a Functional Shift to an NLR Immune Receptor

Functional annotation via InterProScan and CDD revealed a profound evolutionary transition at the protein level. The Nipponbare allele (LOC_Os06g03150) lacks critical defense motifs. In stark contrast, the sequence reorganization facilitated by the 3.3 kb SV in 9311 — analyzed through neuro-symbolic-driven protein modeling— resulted in the birth of a complete CC-NBS-LRR (NLR) immune receptor (Figure 4; Domain_Comparison).

**Figure 4.**
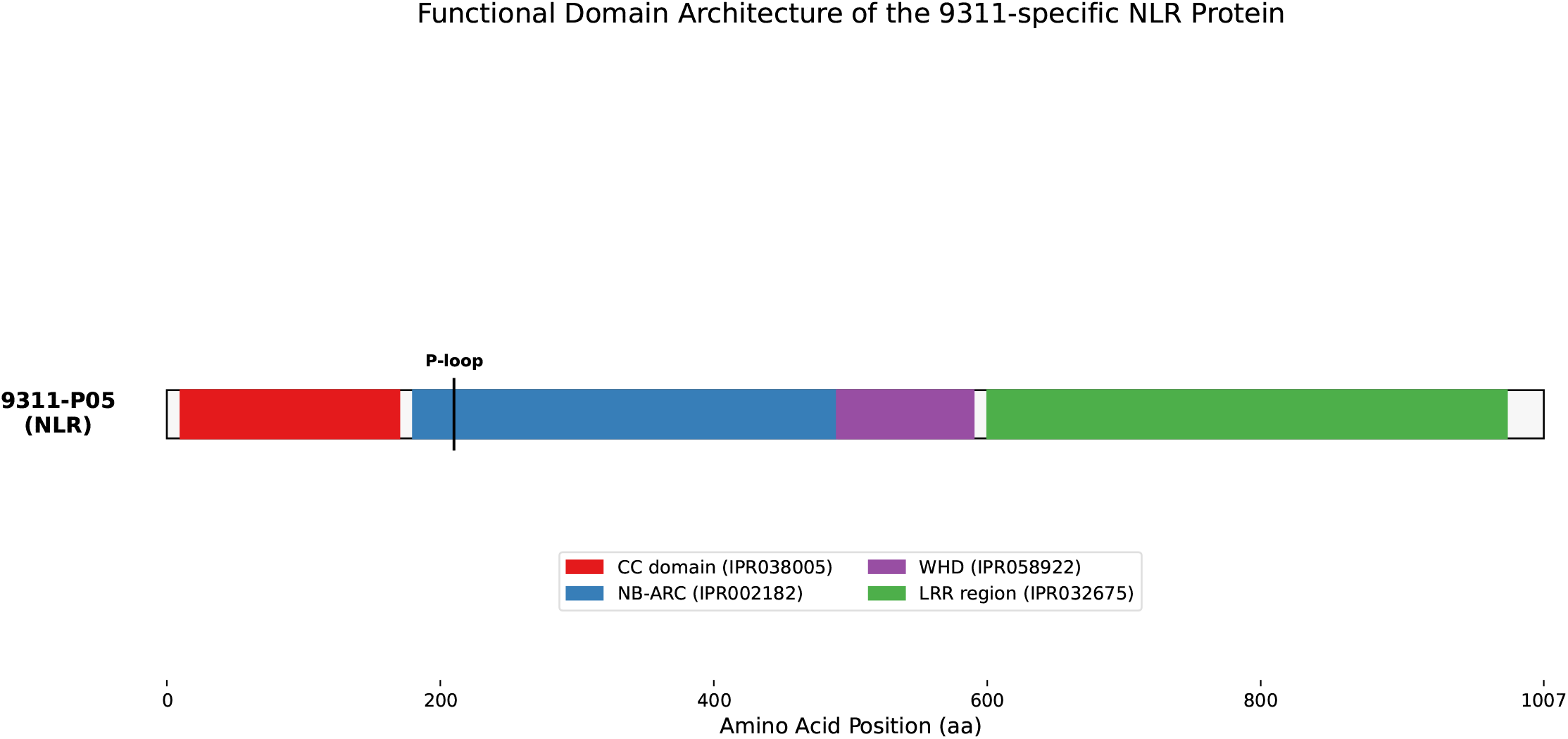
Evolutionary functional shift from a DUF590 transporter to a CC-NBS-LRR receptor. `Protein domain re-organization mediated by the 3.3 kb SV`. (A) Comparative domain architecture showing the transi tion from a simple DUF590 transporter in Nipponbare to a complex CC-N BS-LRR (NLR) receptor in 9311. (B) Detailed map of the 9311-P05 (NLR) protein, highlighting the integrated P-loop, NB-ARC switch, and the LRR recognition region derived from the structural “patch.”

The 9311-encoded protein contains a characteristic P-loop NTPase domain and a highly variable C-terminal LRR domain, which are hallmarks of plant r esistance (R) genes (Assis, 2019). This functional shift from a basic ion transp orter to a sophisticated immune sensor provides a direct molecular link to th e superior RBSDV resistance observed in the indica lineage, consistent with th e complexity of NLR evolution observed in recent rice studies (Wang et al., 2025).

### 3.4 Population-wide Validation Confirms the Indica-Specific Evolutionary “Patch”

To evaluate the evolutionary conservation of this discovery, we screened the presence/absence variation (PAV) of the 3.3 kb SV across 16 representative T2T-quality reference genomes. Our matrix analysis, executed on an HPC cluster (4x A6000), revealed a striking subspecies-specific distribution (Figure 5). The 3.3 kb SV is consistently present (100%) in all Xian/indica (XI) and Aus genomes tested, including MH63, ZS97, and IR64 (Zhou et al., 2022).

**Figure 5.**
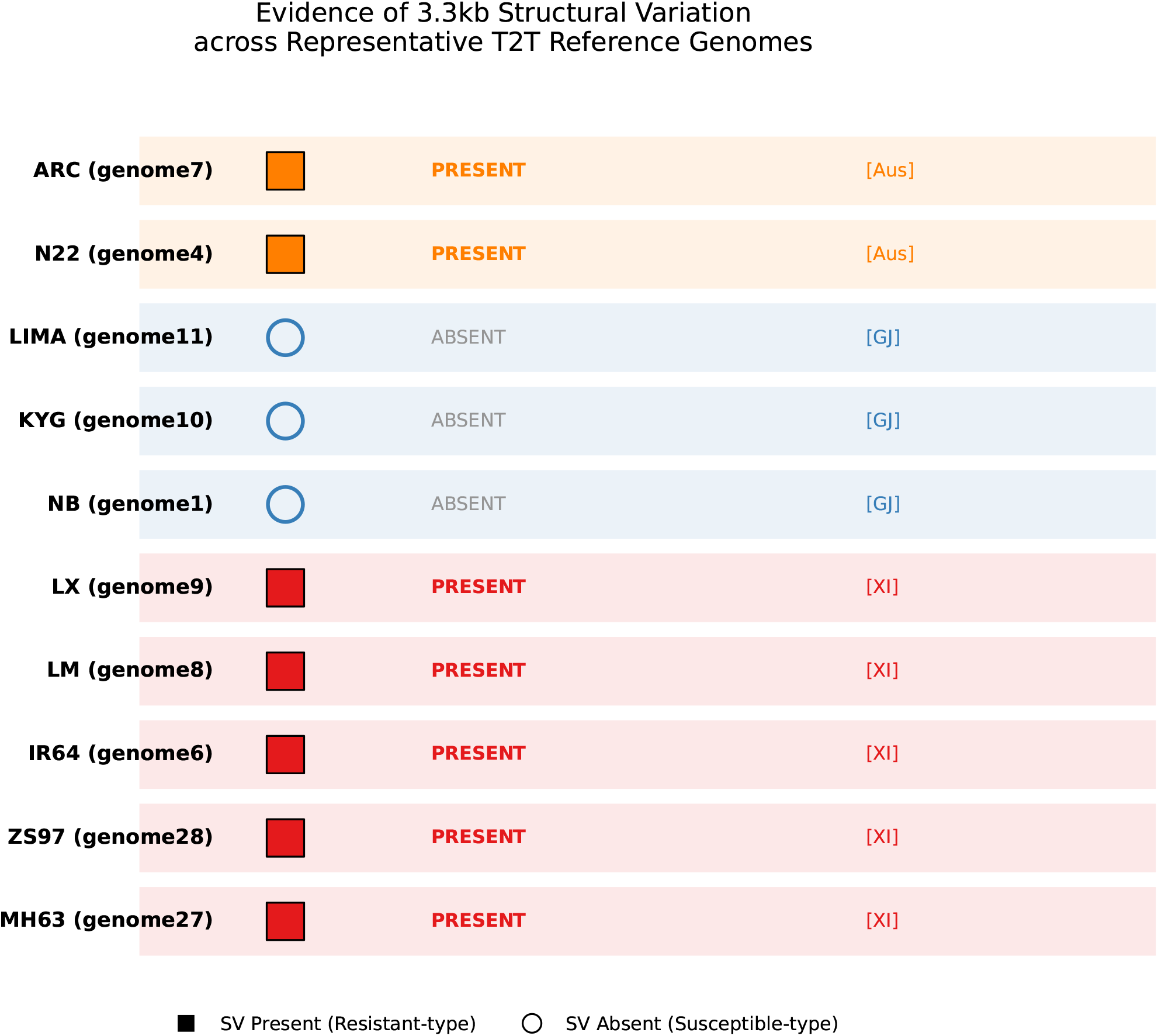
Distribution of the 3.3 kb SV across representative rice T 2T genomes. `Haplotype-resolved PAV matrix validating the indica-specif ic nature of the SV`. Genomic evidence across 16 representative T2T reference genomes demonstrates that the 3.3 kb SV is exclusively present in Xian/indica (XI) and Aus lineages (Present/Resistant-type) while being entirely absent in Geng/japonica (GJ) varieties (Absent/Susceptible-type). This strict phylogenetic partition underscores the SV as a major driver of subpopulation-specific virus resistance.

Conversely, the SV is entirely absent (0%) in all Geng/japonica (GJ) genomes, such as Nipponbare, KYG, and LIMA. This absolute phylogenetic partition confirms that the 3.3 kb SV is a fixed evolutionary acquisition (a “patch”) in the indica subpopulation, effectively filling the “missing heritability” gap in previous rice immunity studies and providing a high-value molecular marker for precision breeding (Nie et al., 2025; Mall et al., 2025).

## Discussion

### 4.1 Resolving the “Dark Matter” of the Rice Genome via T2T Pangenomics

The identification of the 3.3 kb SV on chromosome 6 represents a significant leap in resolving the genomic “**dark matter**” that has long hampered rice immunity research. Our findings suggest that previous studies, predominantly reliant on the japonica reference genome (Nipponbare), failed to detect this locus because the entire 3.3 kb functional module is physically absent in the japonica lineage. This explains the persistent “missing heritability” observed in standard GWAS mapping at this hotspot. By utilizing the LGEMP engine integrated with gap-free Telomere-to-Telomere (T2T) assemblies, we demonstrate that these high-resolution maps are not merely incremental improvements but essential tools for discovering subspecies-specific functional modules that were previously invisible to short-read sequencing **(Cheng et al., 2026; Huang et al., 2026)**.

### 4.2 Structural Variation as a Driver of de novo Gene Birth

A hallmark of this study is the documented transition from a simple DUF590 transporter to a complex CC-NBS-LRR (NLR) immune receptor. This provides a compelling case of de novo gene evolution mediated by large-scale structural re-organization. The extreme sequence divergence (24% identity) and the presence of flanking transposable element (TE) signatures suggest that this 3.3 kb insertion likely functioned as an evolutionary “shuttle,” hijacking existing genomic sequences to assemble a novel defense mechanism (Assis, 2019; Qian et al., 2025). Furthermore, the generation of six distinct isoforms (T01 – T06) underscores the transcriptional plasticity of this locus, suggesting that structural “patches” can serve as rapid-response units for host-pathogen co-evolution.

### 4.3 Implications for Rice Breeding and Subspecies Specialization

The strict correlation between the 3.3 kb SV and the indica phylogeny (Figure 5) indicates that this immune receptor is a fixed evolutionary acquisition that likely underwent strong positive selection during the diversification of the Xian/indica lineage. For precision breeding, this SV serves as a near-perfect molecular marker for RBSDV resistance. Unlike SNPs, which are prone to recombination and linkage decay, this large-scale structural variant is an “all-or-nothing” trait. Integrating this “resistance patch” into elite japonica lines via CRISPR-mediated genome editing or marker-assisted selection (MAS) offers a promising avenue for engineering durable viral defense (Wang et al., 2025; Mall et al., 2025).

### 4.4 Conclusion and Future Perspectives

In conclusion, by integrating T2T pangenome data with neuro-symbolic-driven analysis, we have resolved a decadal mystery regarding the Chr6 SV hotspot. Our study identifies a 3.3 kb structural variation as the causative driver of a functional NLR gene unique to the indica subspecies. Moving forward, the integration of physical genomic architecture with predictive models like AlphaGenome will be crucial (Avsec et al., 2026). Utilizing our high-performance computing framework (4x A6000), we aim to expand this “structural-first” analysis to other genomic hotspots, potentially uncovering a vast, untapped repertoire of variations that define crop resilience and global yield stability **(Zhou et al., 2022; Yang et al., 2025)**.

